# Evidence of multifaceted functions of codon usage in translation within the model beetle *Tribolium castaneum*

**DOI:** 10.1101/754911

**Authors:** Carrie A. Whittle, Arpita Kulkarni, Cassandra G. Extavour

**Affiliations:** Department of Organismic and Evolutionary Biology, Harvard University, 16 Divinity Avenue, Cambridge MA 02138, USA; Department of Molecular and Cellular Biology, Harvard University, 16 Divinity Avenue, Cambridge MA 02138, USA

**Keywords:** Optimal codons, non-optimal codons, translational selection, translation regulation, tRNA genes

## Abstract

Synonymous codon use is non-random. Codons most used in highly transcribed genes, often called optimal codons, typically have high gene counts of matching tRNA genes (tRNA abundance) and promote accurate and/or efficient translation. Non-optimal codons, those least used in highly expressed genes, may also affect translation. In multicellular organisms, codon optimality may vary among tissues. At present however, codon use remains poorly understood in multicellular organisms. Here, we studied codon usage of genes highly transcribed in germ line (testis, ovary) and somatic tissues (gonadectomized males and females) of the beetle *Tribolium castaneum*. The results demonstrate that: 1) the majority of optimal codons were organism-wide, the same in all tissues, and had numerous matching tRNA gene copies (Opt-codon_↑tRNAs_), consistent with translational selection; 2) some optimal codons varied among tissues, suggesting tissue-specific tRNA populations; 3) wobble tRNA were required for translation of certain optimal codons (Opt-codon_wobble_), possibly allowing precise translation and/or protein folding; and 4) remarkably, some non-optimal codons had abundant tRNA genes (Nonopt-codon_↑tRNAs_), and genes using those codons were tightly linked to ribosomal and stress-response functions. Thus, Nonopt-codon_↑tRNAs_ codons may regulate translation of specific genes. Together, the evidence suggests that codon use and tRNA genes regulate multiple translational processes in *T. castaneum*.

## 1. Introduction

In protein coding genes, the synonymous codons of amino acids are not used randomly. Biases in codon usage are thought to result from selection for translational efficiency and/or accuracy.^1–9^ Mutational pressures can also shape codon usage.^5,10–13^ Translational selection in many organisms has been supported by findings that the highly transcribed genes preferentially use a subset of codons, often described as “optimal” codons,^2,6,12–18^ and has been observed in bacteria,^5,6,17^ fungi,^16,19,20^ plants^2,14,21^ and animals, including spiders^22^ and insects (e.g., *Drosophila, Aedes, Anopheles*, *Gryllus*, *Oncopeltus,* and weakly observed in *Bombyx* ^2,15,23–27^). Whole-genome data show that optimal codons typically have correspondingly high numbers of iso-accepting tRNA gene copies in the genome, reflecting an organism’s relative tRNA abundance,^1,5,6,19,20,28^ and is consistent with selection for translational optimization.^1,4,5,18,20,29–33^ The utility of tRNA gene number to quantify organismal tRNA abundance has been supported *in vivo* in bacteria and eukaryotes.^28,34,35^ For instance, the addition of tRNA genes for a codon of a specific amino acid to the *E. coli* genome markedly improved translation rates of genes containing that amino acid.^28^ In this regard, the increased use of optimal codons in highly transcribed genes,^2,5,14^ and the correspondence of these codons to abundant tRNA genes,^1,4^ suggest that selection may favor optimization for cost efficient and/or accurate translation.

In contrast to unicellular systems, in multicellular organisms measuring codon usage can be complicated by the plurality of tissues, as optimal codons and tRNA populations may vary among tissue types.^36-38^ For instance, cellular tRNA abundances can vary among tissues or cell types for at least some codons,^37,39,40^ suggesting that translational selection may differ among tissues.^37^ This has also been supported by findings of some variation in codon use of genes transcribed in different tissues in the few organisms studied to date. For example, in the plant *Arabidopsis* the use of specific codons in a gene depends on the tissue type in which it is maximally expressed, suggesting this species has localized tRNA populations,^38^ a pattern that has also been proposed for rice.^41^ Although similar studies in metazoans have been rare, a recent investigation in *D. melanogaster* showed that codons associated with elevated expression were not universal across tissues. For example, AAT was more commonly used than AAC for Asn in some tissues (e.g., testis, hindgut), while TGT was favored over TGC for Cys in the salivary glands, that was suggested to provide evidence of tissue-specific tRNA populations.^36^ Additional studies are warranted to determine the universality of distinct optimal codon identities in various tissues of an organism. In particular, the germ line and somatic tissues comprise contrasts of significant interest, as the former directly determines an organism’s reproductive success and fitness and experiences haploid selection in the meiotic and sex cells, such that translational optimization may be particularly relevant to those tissues.

While much attention has been focused on optimal codons in the literature, growing experimental research, largely from single-celled models or *in vitro* systems, suggests that non-optimal codons, those codons least used in highly transcribed genes (and/or codons defined as “rare” in some studies), can also play significant regulatory roles in translation.^34,42,43^ In yeast for example, it was shown that cells altered their tRNA populations under stress and had increased levels of tRNAs that matched the rare codons found in stress-response genes, thus allowing the preferential translation of those mRNAs under stressful conditions, without any change in mRNA abundance.^44^ Findings in cyanobacteria have indicated that circadian rhythms are regulated post-transcriptionally based on non-optimized codon use in genes of the *kaiABC1* cluster.^45^ Further, non-optimal codons have been shown to slow rates of translational elongation and to control ribosome traffic on mRNA, which allows proper co-translational protein folding and/or functionality, based on *in vitro* cell-free translation systems from *Neurospora*^7^ and *Drosophila*.^9^ Non-optimal codons have also been found to facilitate co-translational protein folding in various yeast models ^46^ These data show that the use of one or a few types of rare codon(s) in a gene may markedly affect its translation, depending on the tRNA pool, suggesting that the supply-demand relationship between non-optimal codons and their matching tRNA abundances could comprise an adaptive mechanism of translational regulation.^34,44-48^ To further understand this phenomenon, genomics and molecular evolution research on codon usage patterns in animal systems should expand beyond the typical focus on optimal codons, and specifically include assessments of non-optimal codons, and their relationships to tRNA genes.

In addition to non-optimal codons *per se*, some studies have indicated that the use of codons that have no matching tRNA, and obligately require wobble codon-anticodon tRNAs (wobbly at the third nucleotide of the codon) may also influence translation.^34^ For instance, an investigation in four divergent eukaryotes found that the relative translation levels of cell-cycling gene mRNAs during various stages of the cell cycle depended on the frequency of codons that had no corresponding tRNA gene copies in the genome and thus required wobble tRNA.^49^ Further, experimental research in yeast, human cells, and nematodes has shown that obligatory use of wobble tRNA decelerates translational elongation by slowing ribosomal translocation on the mRNA.^34,50,51^ In this regard, the use of codons that require wobble tRNA could have a significant effect on translational dynamics, particularly in slowing translation,^34^ and thus should also be considered in studies of codon usage patterns in an organism.

A metazoan species providing a promising pathway for the comprehensive study of codon usage in a multicellular system is the Coleopteran rust red flour beetle *Tribolium castaneum*. *T. castaneum* is a long standing model for genetics and developmental biology, has a well characterized genome,^18,52,53^ and is estimated to have diverged from the fellow insect *Drosophila* approximately 300 Mya.^54–58^ While a prior pioneering study had identified a putative list of optimal codons for *T. castaneum,*^18^ the approach used in that study involved correlation analyses between codon frequency and expression level. Given that this method has been thought to often be poorly suited to revealing optimal codons, defined as those most common in highly transcribed genes,^1,5,59^ analyses of codon use in this taxon would benefit from being revisited with alternative methods. Optimal codons can be most readily revealed via direct contrasts of codon usage in the highest versus lowest expressed genes in the genome, also known as the contrast method.^2,13–17,21,24,59^ At present, like most multicellular model organisms, a multifaceted integrative approach has not yet been applied to assessments of codon usage in this beetle taxon, including the identification of optimal and non-optimal codons in highly transcribed genes at an organism-wide level, and within the somatic versus germ line tissues, nor have assessments been available of the links been such codon usage and tRNA gene counts, wobble tRNA, and gene functionality.

In the present study, we address these outstanding issues on codon usage in *T. castaneum* using genome-wide protein-sequence datasets (CDS) and large-scale transcriptome datasets from the male and female germ lines and somatic tissues (testes, ovaries, gonadectomized (GT-) males and GT-females).^60^ From these data, we rigorously study optimal and non-optimal codons in this taxon, and their relationships to tRNA abundances and gene ontology. From these analyses, we report strong evidence for organism-wide optimal codons in all four tissue types and both sexes. The majority of these optimal codons have abundant matching tRNAs (Opt_↑tRNA_ status), consistent with pervasive translational selection for efficient and/or accurate protein synthesis in this species. A minority of optimal codons vary among the four tissues, suggesting small, but potentially meaningful, differences in tRNA populations between tissue types. Crucially, we report that a subset of the optimal codons did not have direct tRNA matches and obligately required wobble tRNA for translation (Opt-codon_wobble_), which we propose may comprise a mechanism for slowing translation for accuracy or protein-folding purposes. Finally, we find that a number of non-optimal codons unexpectedly have abundant perfectly matching tRNA gene copies (Nonopt-codon_↑tRNAs_) and that these rare codons are preferentially used in genes with specific functions, including ribosomal protein genes and stress response genes. Thus, we hypothesize that the use of codons with Nonopt-codon_↑tRNAs_ status may be a potential mechanism to ensure preferential translation of specific gene mRNAs. Collectively, our results reveal the multiple roles of codon usage in this beetle, suggesting not just pervasive selection for the use of specific codons in highly transcribed genes for efficient and/or accurate translation, but also translational regulatory roles of wobble codons and of non-optimal codons.

## 2. Materials and Methods

### 2.1. T. castaneum CDS

The annotated CDS of our main target species *T. castaneum* (v.5.2) were downloaded from Ensembl Metazoa (http://metazoa.ensembl.org) and are also available at BeetleBase^52,53^). The full CDS per gene (longest CDS per gene, N=16,434) was used for the study of codon usage. The full genome and its descriptive GFF file was also downloaded for assessments.

### 2.2. Biological samples and RNA-seq

We aimed to determine the expression level (FPKM) for each of 16,434 genes in *T. castaneum* for germ line and somatic tissues. For this we used the large-scale RNA-seq datasets for the ovaries, testes, GT-females and GT-males shown in Supplementary Table S1.^60^ The *T. castaneum* specimens were provided by the Brown lab at KSU (https://www.k-state.edu/biology/people/tenure/brown/). Samples were grown under standard conditions until adulthood and tissue dissections were then performed on unmated adults (a total of 150 animals per sex per biological replicate), and RNA was extracted and processed for RNA-seq, as described previously.^60^

### 2.3. Gene expression

The RNA-seq reads (76bp) per sample were trimmed of adapters and poor-quality bases using the program BBduk available from the Joint Genome Initiative (https://jgi.doe.gov/data-and-tools/bbtools/) set at default parameters.

Gene expression level was determined for the 16,434 genes (CDS) as FPKM after mapping each RNA-seq dataset per tissue to the full CDS list for each species using Geneious Read Mapper^61^, which yielded highly similar results as other common mappers such as BBmap (https://jgi.doe.gov/data-and-tools/bbtools/). The average FPKM across samples per tissue type (Supplementary Table S1) was used to measure expression per tissue. FPKM values were highly correlated between replicates of each sample type (Spearman’s Ranked R>0.9, P<2×10^−7^)

### 2.4. Identification of optimal and non-optimal Codons

For identification of the optimal codons, we measured the relative synonymous codon usage (RSCU) per codon per amino acid for each gene under study using CAICal.^62^ RSCU values indicate the relative usage of a codon in a synonymous codon family, and values >1 and <1 indicate favored and unfavored usage as compared to that expected under equal usage of all codons respectively, and greater relative RSCU values among codons indicates elevated usage. For each of the 18 amino acids in the genetic code with synonymous codons (note that Trp and Met only have one codon each), we identified the optimal codon using the contrast method.^13–15,17,21,24,59,63^ For this, we determined the difference in RSCU (ΔRSCU) per codon between genes with the highest 5% versus the lowest 5% expression. The primary optimal codon for each amino acid was defined as the codon with the highest and statistically significant positive ΔRSCU value, indicating preferred usage in highly transcribed genes.^13–15,17,21,24,59,63^ The primary non-optimal codon per amino acid was defined as the codon with the largest negative and statistically significant ΔRSCU value, indicating low usage in highly transcribed genes. Statistical significance per codon was applied using a t-test between RSCU values across all genes for high versus low expressed genes.

As the literature reflects some variation in codon use terminology among studies to date, we explicitly define the term “optimal codons” herein as those codons most used in highly transcribed genes based on ΔRSCU, which infers an innate advantage of the codon under high transcription. Then, we secondarily assessed each optimal codon’s correspondence to the number of matching (codon-anticodon) tRNA genes in order to test their role in translational accuracy/efficiency^1,4,5,18,20,29–33^ or to infer possible other functions (e.g., wobble codons for translational slowing). For non-optimal codons a similar approach was used wherein the non-optimal codon status was identified based solely on ΔRSCU, and their relationships to tRNA were then separately assessed.

The frequency of optimal codons (Fop) is a measure of the degree of optimal codon usage per gene.^6^ Fop was determined in CodonW^64^ using the primary optimal codons identified herein. Fop was also determined using the primary optimal codons previously identified by Williford and Demuth 2012.^18^ As multiple codons per amino acid were classified as optimal in that assessment, we defined each primary optimal codon from the study as that with the strongest average positive correlation across tissues for measuring Fop.

For an additional layer stringency, we wished to exclude the possibility that expression-mediated mutational-biases towards specific nucleotides, which have been observed to some extent in certain organisms to date (e.g., *E. coli*, humans^65,66^), contribute towards codon differences among high and low expressed genes herein. For this, we extracted all introns for every gene in the genome (those with introns) using the GFF file available (see section 2.1). Introns are thought to be mostly selectively neutral,^18,67^ and thus the nucleotide content should reflect any underlying mutational pressures in the genome, and on the nucleotide composition of synonymous codons in an organism.^13,18,67^ If mutational pressures on introns are not associated with gene expression level, it will exclude this factor in causing optimal codons in the highly expressed genes, and further affirm the role of selection. All introns that were >50bp were extracted as the region between exons and were concatenated per gene. The association between GC content and expression level were assessed using a scatter plot and Spearman’s ranked R.

### 2.5. Identification of tRNA Genes

To assess whether or how the optimal and non-optimal codons were related to the tRNA gene copy number, we determined the number of iso-accepting tRNA genes per codon in the genome (*T. castaneum* v. 5.2) using tRNA-scan SE.^18,53,68^ The list of tRNA gene numbers identified in the current genome version was identical to that reported previously^18^ and is shown in Table 1.

**Table 1.**
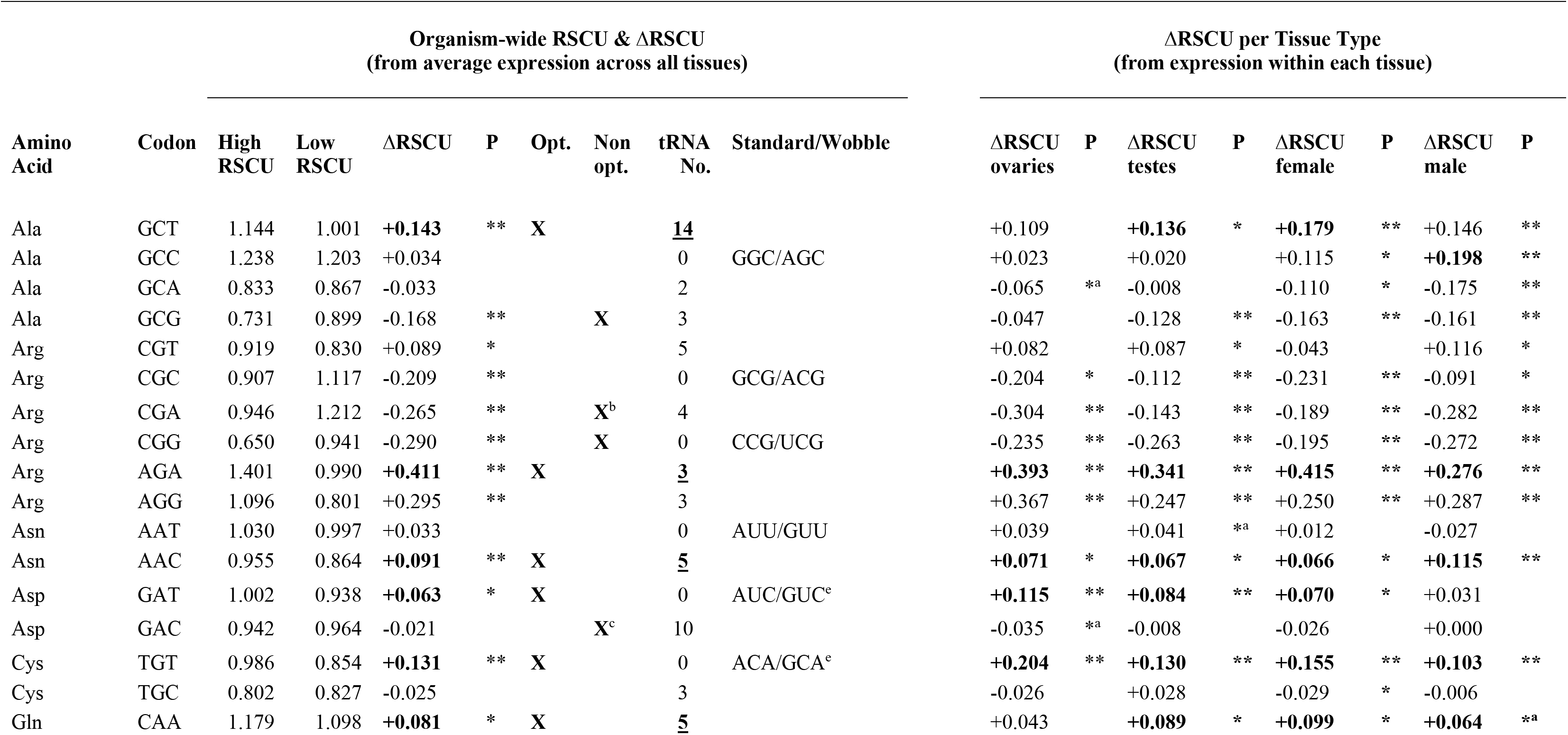
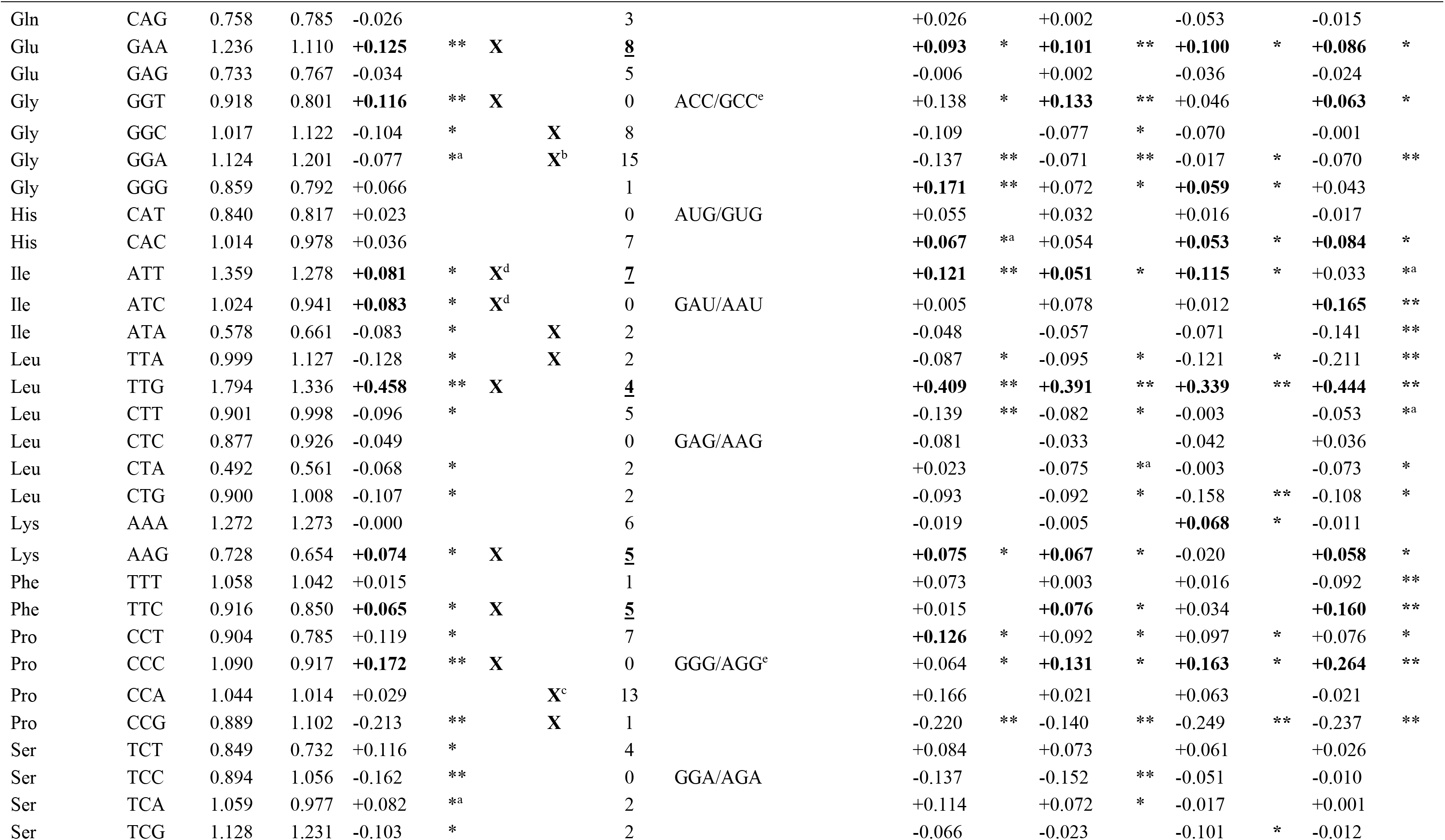
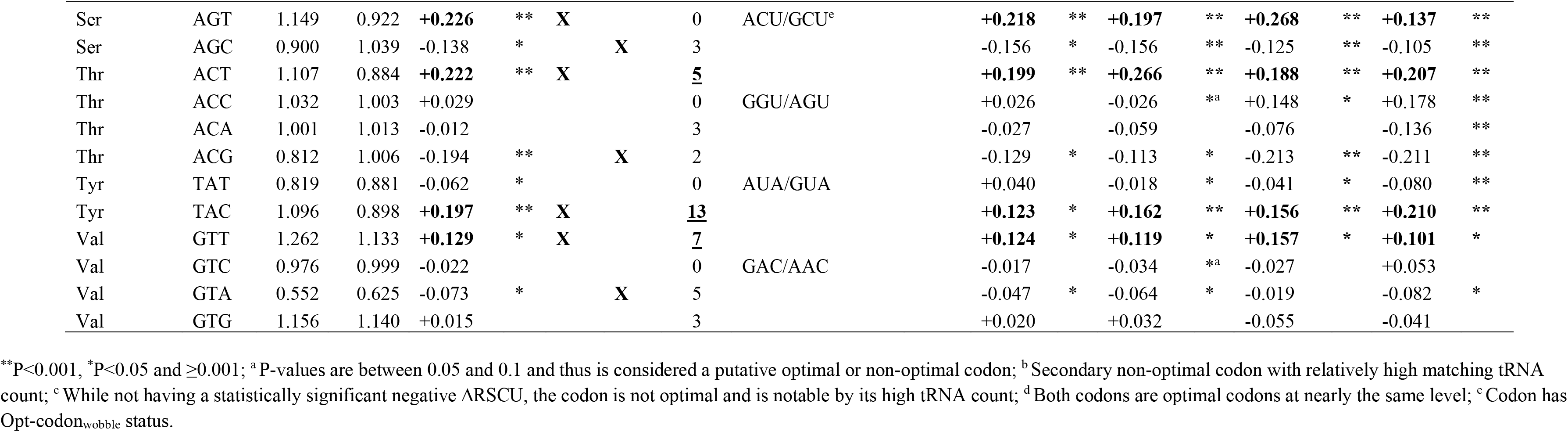
The organism-wide ΔRSCU between high versus low expressed genes (using averaged expression across all four tissue types, the ovaries, testes, GT-females, and GT-males). In addition, the ΔRSCU are shown when high and low expressed genes were determined for each of the four individual tissue types. The primary optimal (Opt.) codons are in bold and have the largest positive and statistically significant ΔRSCU (t-test P<0.05) per amino acid. For the combined four tissue assessment (organism-wide), the primary optimal (Opt.) and non-optimal codons (Non opt.) are shown with X. Cases where relatively plentiful tRNA genes match the optimal codon per amino acid are underlined and bold. The wobble anticodons for codons with zero matching tRNA copies are shown (standard anticodon/wobble anticodon shown according to classical wobble rules; see also^81,94^).

### 2.6. GO Functions

The predicted GO functions were determined using Panther^69^ using the option for *T. castaneum* as species

### 2.7. Data Availability

The CDS and genome v. 5.2 for *T. castaneum* are available at Ensembl Metazoa (http://metazoa.ensembl.org). RNA-seq data for all samples from *T. castaneum* described in Supplementary Table S1 are available at the SRA database under Bio-project number PRJNA564136.

## 3. Results and Discussion

### 3.1. Optimal codons in *T. castaneum*

We first report the organism-wide, or global, optimal codon per amino acid for *T. castaneum* using ΔRSCU and the average expression levels of all annotated genes across all four studied tissue types (testis, ovary, GT-male, GT-female) in Table 1. The primary optimal codon was defined as the codon with the largest positive ΔRSCU between highly and lowly transcribed genes and with P<0.05), was found for 17 of the 18 amino acids with synonymous codons. Seven primary optimal codons ended in T, three in A, five in C and two in G. We noted that Ile had two codons with nearly identical ΔRSCU values. Further, CAC for His showed signs of optimal codon usage in several individual tissues (see following section), and including this codon yields a study-wide total of 18 optimal codons (Table 1). The range of ΔRSCU values is similar to or larger than that observed in other multicellular eukaryotes, including nematode species*, Drosophila, Populus* and *Neurospora.* ^2,14–16^ Thus, the patterns in Table 1 are consistent with selection pressures have favored the use of a specific subset of codons in highly expressed genes^5^ (for results on non-optimal codons see section 3.4 below).

While the striking use of specific optimal codons in genes under high expression levels in Table 1 in itself provides evidence of selection on codon usage, we wished to include additional layers of stringency to affirm the role of selection in favoring these codons. First, we determined the frequency of optimal codons (Fop), a measure of the degree of optimal codon usage per gene,^6^ for all studied genes in the genome (N=16,434). As shown in Fig. 1A, we found that the Fop increased from genes with low (top 5% in the genome), to moderate (5 to 95%), to high (top 5%) expression levels (Ranked ANOVA and Dunn’s paired test P<0.05). As low and high expressed genes were used to identify the optimal codons, the Fop was expectedly lowest and highest in those categories of genes respectively. Importantly however, moderately expressed genes, which were not used to identify the optimal codons, showed intermediate Fop values, suggesting a genome-wide tendency for greater use of optimal codons in CDS with elevated expression. Second, as codon usage can vary with protein length in some eukaryotes,^2,5,70^ we repeated the assessment in Fig. 1A using genes with similar CDS lengths, which we binned into short (<150 codons), medium (≥150, <300), and long CDS (≥300). For each of these three length categories, we found the same stepwise increase of Fop values with expression level (Ranked-ANOVAs P<0.001). Thus, the link between expression and optimal codons cannot be explained by protein length. Third, from examination of introns, wherein nucleotide content is mostly shaped by mutational pressures,^18,67,71^ we found that the GC (and thus AT) content of introns was uncorrelated to gene expression level (Spearman’s correlation R= −0.09, Fig. 1B),^72,73^ and thus indicates an absence of expression-mediated mutational biases^12,65,66,71^ in this species. Further to this point, unlike some organisms wherein optimal codons typically end in only two or three types of nucleotides,^2,14,21,24^ all four nucleotides are represented at the terminal position of optimal codons of this species (Table 1); this also excludes mutational biases in shaping the optimal codons in highly transcribed genes in this taxon.^5^ Taken together, while we do not exclude the possibility that non-selective (mutational) mechanisms may contribute toward codon use of genes, particularly those under low or even moderate expression,^74^ our observations indicate that a history of selection pressures likely plays a significant role in shaping the codon use of the most highly transcribed genes in this organism (top 5% expression), shown in Table 1.

**Figure 1.**
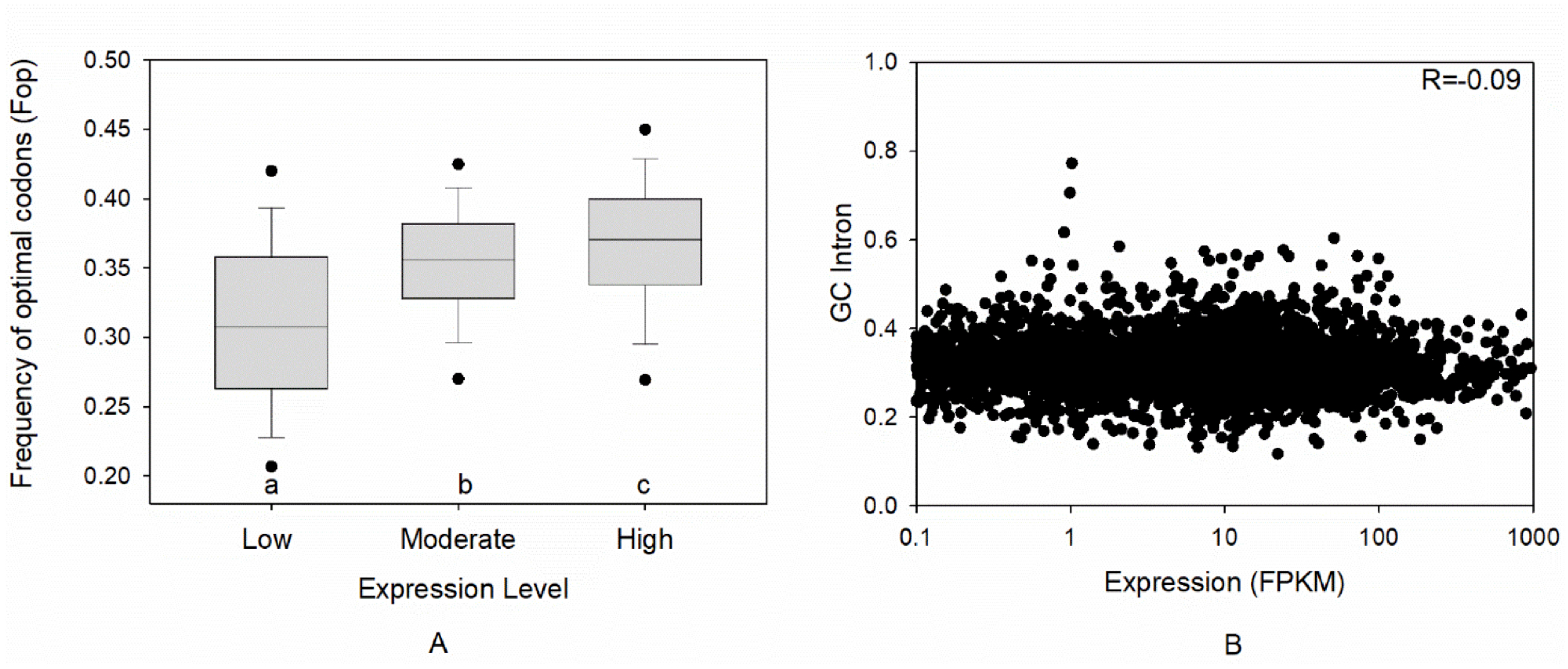
**A.**The frequency of optimal codons (Fop) across all 16,434 genes studied in *T. castaneum*. Genes are categorized into low (lowest 5%, FPKM<0.013), moderate (5 to 95%) and high (top 5%) transcription (FPKM>103) groups based on average expression across all four tissue types (testes, ovaries, GT-males, GT-females). Different letters below bars indicate a statistically significant difference using Ranked ANOVA and Dunn’s paired contrasts (P<0.05). **B.**The GC content of introns with respect to the expression level per gene (Spearman’s Ranked R is shown). Values are shown for all genes with introns >50bp (N=5,143).

### 3.2. Most, but not all, optimal codons are the same across germ line and somatic tissues

In order to compare optimal codon usage among the tissues under study, we next determined the optimal codons (using ΔRSCU) using genes with high versus low expression (top and lowest 5%) separately for each of the four individual tissue types, ovaries, testes, GT-females and GT-males. For rigor in this assessment, we identified the subset of genes in the top 5% expression class that were only in the top category for one tissue type (and were not in the top 5% expression in any of the other three tissues), to discern whether or not there was a tissue effect on optimal codons. Under these criteria, we identified 372, 450, 444, and 272 genes for analysis, for ovaries, testes, GT-females and GT-males respectively. This allowed us to specifically assess the codon usage of genes that were maximally transcribed only in one individual tissue, as it has been found that if tissue-type has an effect on codon use, this effect is most apt to be evident in its highly transcribed genes^38^. The results for ΔRSCU per tissue type are shown in Table 1. We report that 15 of the 18 primary optimal codons (including His) from the organism-wide assessment were identified as having the same optimal codon in three, or all four, of the individual tissue-types (Table 1). Thus, the vast majority of primary optimal codons were the same in these divergent tissues, including male and female germ lines and somatic tissue types.

However, several significant differences were also observed among tissues. For example, a male-specific primary optimal codon was identified for the amino acid Phe (with two synonymous codons), as the codon TTC was optimal in the testes and GT-males, but not in the ovaries or GT-females (Table 1). Similarly, a GT-male-specific primary optimal codon ATC was identified for Ile (with three synonymous codons), where ATT was optimal for the other three tissues. In turn, an ovary-specific optimal codon was evident for Pro (with four synonymous codons), as the primary optimal codon was CCC in all tissues except for the ovaries, where it was CCT. In addition, a GT-female optimal codon was identified for Lys (two synonymous codons), where AAG was optimal in the ovaries, testes, and GT-males, but its alternate codon AAA was optimal for GT-females. These examples show that the primary optimal codon varies among tissue types in this beetle, and thus this pattern suggests that translational selection regimes, and thus corresponding tRNA populations may also vary among tissues.^36^ Further, it is worth noting that in some cases there may be tissue-specific preferences for codons using wobble tRNA (e.g., ATC for Ile in GT-males, see section 3.3).

These present results are consistent with the few available studies of tissue-specific codon usages and translational selection from the fellow insect *D. melanogaster*^36^ and in studied plants^38,41^ (note that although some evidence suggests humans have tissue-specific optimal codons, this has been debated, and may largely be an effect of the GC content of isochores, which exist in those organisms^75,76^). Together, while the vast majority of optimal codons are shared across tissues in these beetles, non-negligible differences are observed between tissues and sexes. Direct quantification of tRNAs in cells or tissues has been mostly restricted to date to lab models of bacteria, yeast or *in vitro* human cell lines,^37,39,40,44,77^ and the accuracy and limitations of the various approaches (based on microarrays, Northern blot, quantitative PCR, RNA-seq) remains debated^40,44,78,79^. Nevertheless, the development of robust methods to sequence tRNAs that are applicable to non-traditional model organisms will allow further tests of whether or how tRNA expression levels vary with tissues in *T. castaneum*, as is strongly suggested by these results.^36^

### 3.3. A majority of organism-wide optimal codons have high tRNA gene copy numbers

Given the minimal differences among tissues, for our remaining analyses we focus on the organism-wide optimal codon usages (Table 1). The number of tRNA gene copies in the genome has commonly been used as a measure of the relative abundance of each tRNA species.^1,4,18,20,29,30,49^ If optimal codon usage were consistently a result of selection in response to abundant tRNAs, then the primary optimal codon per amino acid should also have high relative tRNA gene frequency (Opt-codon_↑tRNAs_ status). When using the organism-wide optimal codon list (Table 1), we found that 12 of the primary optimal codons also had the highest, or near the highest tRNA gene counts of all codons per amino acid, GCT (Ala), AGA (Arg), AAC (Asn), CAA (Gln), GAA (Glu), ATT (Ile), TTG (Leu), AAG (Lys), TTC (Phe), ACT (Thr), TAC (Tyr), and GTT (Val). Further, while the positive ΔRSCU of CAC for His was not statistically significant using the organism-wide assessment (P=0.26), this codon was optimal when individually considered in the ovaries, GT-females and GT-males (P<0.05), and had seven matching tRNA genes. Thus, when including CAC for His as a codon with optimal status, yields a study-wide total of 13 of the 18 primary optimal codons that have plentiful matching tRNA genes. In other words, a majority of optimal codons have Opt_↑tRNA_ status. These results strongly suggest translational selection for accuracy and/or efficiency^1,4^ across a majority of amino acids in this beetle.

#### Hypothesis 1: Optimal codons use wobble tRNA to resolve conflict of high translation with sequence fidelity

While 13 optimal codons had a high number of direct tRNA matches as expected under selection for optimization of efficient and accurate translation, for the remaining five amino acids, a much different pattern was observed. Specifically, the primary optimal codon (highly used in abundant transcripts) had no direct matching tRNA-genes, and a wobble tRNA (shown in Table 1) must thus be employed for translation of these codons (denoted as Opt-codon_wobble_). For instance, Opt-codon_wobble_ status was observed for the amino acids Asp (GAT), Cys (TGT), Gly (GGT), Pro (CCC) and Ser (AGT). Thus, this result shows that while these identified optimal codons are preferred in highly transcribed genes, their innate benefit cannot be due to having abundant direct matching tRNA, and thus another mechanism must explain their high usage. Further, as shown in Supplementary Text File 1 and Fig. S1, within the group of highly transcribed genes, each of these five individual codons with Opt-codon_wobble_ status showed strong associations with protein length, inferring putatively significant roles of the use of these types of codons in the translation of abundant mRNAs, which may vary with the length of the translated sequence.

Experimental studies in bacteria and eukaryotic models have shown that codons using wobble tRNA act to slow translation by decelerating the translocation of ribosomes on mRNA.^34,50,51^ In addition, a study of the genomes of various eukaryotes (humans, yeast, *Arabidopsis*) have indicated that cell-cycle genes had high usage of codons that had no matching tRNA genes in the genome, and thus must employ wobble tRNA, which inherently have lower codon-anticodon binding affinity than those codons with perfect matches.^49^ The differential use of codons using wobble tRNA in cell-cycle genes, combined with potential oscillations in tRNA abundances, were proposed to differentially regulate the translation rates of gene mRNAs during various stages of the cell cycle.^49^ Further, this was speculated to possibly comprise a broader evolutionarily conserved phenomenon for translational regulation in eukaryotes.^49^ In addition, the usage of wobble-tRNAs in a gene could have some parallel functions to the use of non-optimal codons with low tRNA abundance (Nonopt-codon_↓tRNAs_; see Table 1 for Nonoptimal codons with few tRNAs) which can prevent jamming of multiple ribosomes during the initiation of translation,^35^ and/or slow or pause translation during elongation, which would facilitate accurate protein-folding.^7,9,39,80^ In this regard, the results from these various studies suggest that the slowing of translation that is induced by wobble-tRNA^34,50,51^ could comprise an evolutionarily conserved mechanism shaping various aspects of translation.

Significantly, a key modification that mediates wobbling at the first anticodon position (position 34 of the anticodon loop) is for A34, which may be enzymatically deaminated by adenosine deaminase tRNA (ADATs) to form inosine (I34). The I34 can pair with mRNA 3’codon bases A, C, or U in Eukarya ^81,82^ (see also for an A37 ADAT (*Adat1*) in *D. melanogaster*^83^). For A34 modifications in eukaryotes, available research to date suggests that deamination requires the ADAT2/ADAT3 (hetADAT) enzymes, which are thought to allow A34 modifications across diverse eukaryotic systems. ^81,84^ This modification would be essential for some codons obligately requiring wobble tRNA (those with no matching tRNAs, and no matching unmodified wobble tRNAs) in the highly transcribed genes studied here, including with Opt-codon_wobble_ status (e.g., Pro, CCC, Table 1). Thus, in addition to wobble codons using unmodified tRNAs, further functional study of ADATs is warranted in model insects, such as *T. castaneum*, including possible variation in expression and activity among tissues, in order to help to further ascertain the potential consequences of use of wobble codons requiring tRNA modification at A34 on translation rates and protein folding.^42,81,84^

Taken together, we hypothesize here that for this beetle, the use of codons with Opt-codon_wobble_ status in highly expressed genes comprises a mechanism to slow or pause translation at various sites, which may lead to increased accuracy of translation or allow co-translational protein folding,^50^. In addition, the high frequency of five specific codons with Opt-codon_wobble_ status in genes with abundant mRNAs (Table 1), suggests that these codons might also play a significant role in post-transcriptional differential regulation of protein levels^49^ in these beetles. Additional studies of protein levels of genes with high usage of codons with Opt-codon_wobble_ status will be needed to further test this aspect of the hypothesis.

### 3.4. Certain non-optimal codons have abundant tRNA genes

Herein, we defined the primary non-optimal codon per amino acid stringently as the codon with the largest negative ΔRSCU per amino acid, rather than simply all codons that were not optimal. Using these data, we assessed whether those codons with low usage in highly transcribed genes also exhibit few tRNA gene copies, as might be expected if codon usage is mostly shaped by translational selection for efficient and accurate translation (i.e., for adaptation of optimal codons and tRNA abundance). The organism-wide primary non-optimal codons (per amino acid) are shown in Table 1.

The results showed that some non-optimal codons, as expected, had low numbers of matching tRNA genes (Nonopt-codon_↓tRNAs_ status, e.g., two tRNA genes for ACG (Thr), ATA (Ile), and TTA (Leu), one for CCG (Pro)). Unexpectedly, however, certain non-optimal codons had relatively moderate to high tRNA gene abundance (denoted as Nonopt-codon_↑tRNAs_). For instance, for Arg, whilst the codon CGG had no tRNA gene copies, its sister non-optimal codon CGA (ΔRSCU= −0.290 and −0.265 respectively) had four tRNA gene matches. For Gly, both the primary and secondary non-optimal codons GGC and GGA (−0.104 and −0.077 respectively) had eight and 15 matching tRNA gene copies respectively. For Val, the primary non-optimal codon GTA had five tRNA genes, only slightly lower than the seven observed for its optimal codon GTT. We noted, that if we relaxed our definition of a non-optimal codon to consider any codon that is not optimal, we found that some of those codons also had many corresponding tRNA genes. For example, for Pro the non-optimal codon CCA (which had a weak and nonsignificant positive ΔRSCU value, +0.029, and thus would not have satisfied our strict definition of having the largest negative ΔRSCU for this amino acid) had 13 tRNA genes, an extraordinarily high value compared with other codons. Moreover, for Asp, the (less stringently) defined non-optimal codon GAC had ten matching tRNA copies. Collectively, it is evident that codons that are not the optimal codons in this taxon are not inevitably linked to a low abundance of matching tRNA genes, and rather in some cases exhibit high matching tRNA gene counts. Thus, these patterns suggest it is possible that non-optimal codons with elevated tRNAs play a specific regulatory role for highly transcribed genes.

A recent study in yeast has indicated that stress genes may preferentially use non-optimal codons that have abundant iso-accepting tRNA genes, to increase effective gene expression by promoting their translation over other proteins rather than affecting mRNA levels.^44^ Based on this notion, we hypothesize here that codons with Nonopt-codon_↑tRNAs_ status in *T. castaneum* may regulate the translation of abundant mRNAs of proteins with specific functions in this beetle. To further evaluate this possibility, we examined the predicted gene ontology functions of the highly transcribed genes that had relatively elevated usage of non-optimal codons with abundant matching tRNAs.

#### Hypothesis 2: Non-optimal codons post-transcriptionally regulate translation based on protein functions

We assessed the GO functions of highly transcribed genes (top 5% in the genome from the organism-wide analyses across all four tissues, N=822; and a cutoff of 103.3 FPKM) that had relatively elevated use of codons with Nonopt-codon_↑tRNAs_ status (Table 1). For this assessment, rather than assess all strictly defined non-optimal codons, we chose as examples the codons GGC for Gly, GTA for Val, and CGA for Arg. These three codons were defined as non-optimal by our strict definition (having a large negative and statistically significant ΔRSCU, Table 1) and had substantial matching tRNA gene copy counts (four to eight tRNA genes each). These codons also had negative ΔRSCU values in all four of the tissue types studied (Table 1), indicating they consistently have non-favored status in this organism.

For the amino acid Gly, we identified those highly transcribed genes that had RSCU values for GGC of >1.5. An RSCU value of one is expected for each of the four Gly codons under equal usage, and thus values of 1.5 to 4 for GGC are relatively high. Thus, while Nonopt-codon_↑tRNAs_ are by definition rare in highly expressed genes, this approach allowed us to specifically examine the functions of this group (of highly expressed genes) that had unusually elevated use (RSCU) of this codon with Nonopt-codon_↑tRNAs_ status. A total of 20.4% of the highly transcribed gene set was in this class. As shown in Table 2, these genes included those involved in oxidative stress response, such as Peroxiredoxin, and those involved in olfactory activity. Thus, we speculate that these types of genes, which use codons with Nonopt-codon_↑tRNAs_ status, will exhibit less tRNA competition during translation elongation than those genes that use codons with few or no matching tRNA genes, such as the fellow Gly codon GGG (with only one tRNA match), or even those genes using non-optimal codons for other amino acids, such as CCG for Pro (with one tRNA match) (Table 1). In addition, we found that genes with elevated GGC frequency encoded numerous (N=15) ribosomal proteins. Thus, this finding suggests that usage of the non-optimal codon GGC may shape translation via a second mechanism: namely, by shaping the cellular abundance of specific ribosomal proteins *per se*, which are needed for translation. In this regard, the non-optimal codon usage profiles in Gly appear consistent with a hypothesis wherein the usage of GGC regulates the translation of a subset of genes in this taxon, and may even regulate translation rates *per se* via effects on certain ribosomal proteins.

**Table 2.**
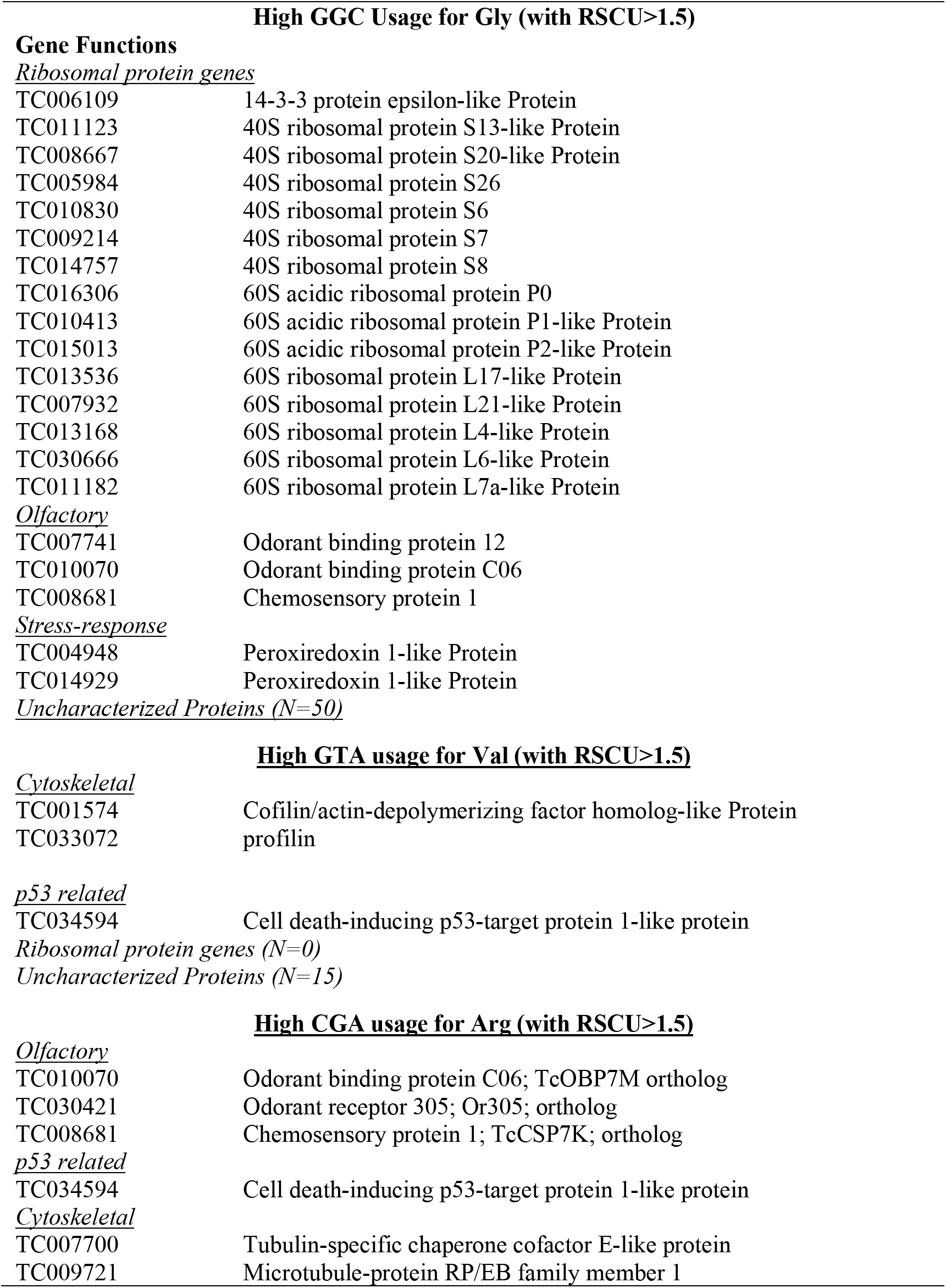
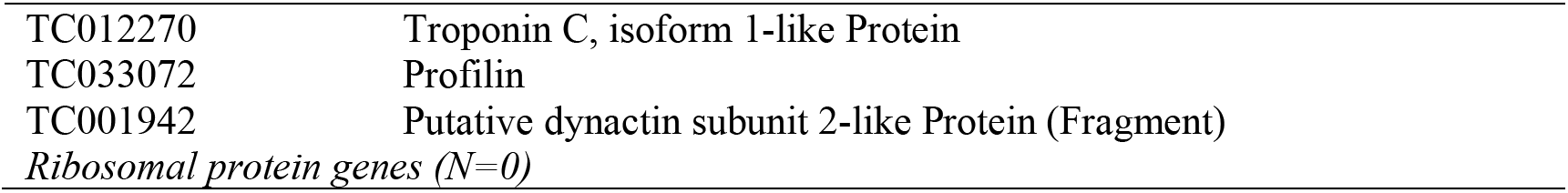
Examples of functions of the highly transcribed genes in *T. castaneum* that have elevated use of codons with Nonopt-codon_↑tRNAs_ status (non-optimal codons with abundant matching tRNA genes ((≥4)). While these codons are by definition typically uncommon in highly transcribed genes (Table 1), the subset of genes with elevated use of these codons, RSCU >1.5, were identified and are shown below. These genes are candidates for translational upregulation due to the elevated use of codons with Nonopt-codon_↑tRNAs_ status.

| **High GGC Usage for Gly (with RSCU>1.5)** |  |
| --- | --- |
| **Gene Functions** |  |
| <u>Ribosomal protein genes</u> |  |
| TC006109 | 14-3-3 protein epsilon-like Protein |
| TC011123 | 40S ribosomal protein S13-like Protein |
| TC008667 | 40S ribosomal protein S20-like Protein |
| TC005984 | 40S ribosomal protein S26 |
| TC010830 | 40S ribosomal protein S6 |
| TC009214 | 40S ribosomal protein S7 |
| TC014757 | 40S ribosomal protein S8 |
| TC016306 | 60S acidic ribosomal protein P0 |
| TC010413 | 60S acidic ribosomal protein P1-like Protein |
| TC015013 | 60S acidic ribosomal protein P2-like Protein |
| TC013536 | 60S ribosomal protein L17-like Protein |
| TC007932 | 60S ribosomal protein L21-like Protein |
| TC013168 | 60S ribosomal protein L4-like Protein |
| TC030666 | 60S ribosomal protein L6-like Protein |
| TC011182 | 60S ribosomal protein L7a-like Protein |
| <u>Olfactory</u> |  |
| TC007741 | Odorant binding protein 12 |
| TC010070 | Odorant binding protein C06 |
| TC008681 | Chemosensory protein 1 |
| <u>Stress-response</u> |  |
| TC004948 | Peroxiredoxin 1-like Protein |
| TC014929 | Peroxiredoxin 1-like Protein |
| <u>Uncharacterized Proteins (N=50)</u> |  |
| **High GTA usage for Val (with RSCU>1.5)** |  |
| <u>Cytoskeletal</u> |  |
| TC001574 | Cofilin/actin-depolymerizing factor homolog-like Protein |
| TC033072 | profilin |
| <u>p53 related</u> |  |
| TC034594 | Cell death-inducing p53-target protein 1-like protein |
| <u>Ribosomal protein genes (N=0)</u> |  |
| <u>Uncharacterized Proteins (N=15)</u> |  |
| **High CGA usage for Arg (with RSCU>1.5)** |  |
| <u>Olfactory</u> |  |
| TC010070 | Odorant binding protein C06; TcOBP7M ortholog |
| TC030421 | Odorant receptor 305; Or305; ortholog |
| TC008681 | Chemosensory protein 1; TcCSP7K; ortholog |
| <u>p53 related</u> |  |
| TC034594 | Cell death-inducing p53-target protein 1-like protein |
| <u>Cytoskeletal</u> |  |
| TC007700 | Tubulin-specific chaperone cofactor E-like protein |
| TC009721 | Microtubule-protein RP/EB family member 1 |
| TC012270 | Troponin C, isoform 1-like Protein |
| TC033072 | Profilin |
| TC001942 | Putative dynactin subunit 2-like Protein (Fragment) |
| *Ribosomal protein genes (N=0)* |  |

In terms of Val, those genes with high expression (top 5% in the genome), very rarely used the identified primary non-optimal codon GTA. In fact, only 5.1% of the 822 highly transcribed genes had GTA RSCU values >1.5, an extraordinarily low frequency. Those that did exhibit RSCU values >1.5 included genes involved in cytoskeleton functions and actin synthesis, such as Cofilin/actin-depolymerizing factor homolog-like protein and profilin, as well as a p53-related cell death protein, and a number of uncharacterized proteins (Table 2). For Arg, which has six synonymous codons, genes with RSCU values >1.5 for the non-optimal codon CGA included genes involved in olfactory signaling and with cytoskeleton roles (Table 2). It is particularly noteworthy that unlike the genes with elevated RSCU for GGC (Gly), which included abundant ribosomal protein genes, no ribosomal protein genes were among those with elevated frequency of GTA in Val or CGA for Arg. Thus, the ribosomal proteins in particular appear to be strongly connected to the usage of the non-optimal GGC Gly codon, and thus we speculate that this codon may be particularly essential to their regulation.

As mentioned above, prior data have suggested that non-optimal codons, when combined with low tRNA abundance, can play important regulatory roles by preventing the jamming of multiple ribosomes during initiation of translation, or slowing translation elongation and facilitating precise protein-folding.^7,9,35,39,80^ The present study, however, shows an additional, and much different, plausible effect of non-optimal codons in *T. castaneum*. Specifically, we show that the use of non-optimal codons with abundant tRNA genes (Nonopt-codon_↑tRNAs_) is tightly linked to predicted gene functionality (Table 2), and thus these codons may be likely to contribute to the preferential translation of mRNAs of specific types of genes. This notion agrees with recent experimental data in yeast suggesting that non-optimal or rare codons in stress genes promote the preferential translation of their mRNA in cells in response to stress-induced changes in tRNA pools.^44^ Thus, this comprises a potential mechanism for preferential translation of specific mRNA. Herein, however, given that abundant tRNA gene copies are available in the genome for codons with Nonopt-codon_↑tRNAs_ status (and thus tRNAs should be consistently abundant in cells), we speculate that the use of these non-optimal codons in certain ribosomal protein and stress genes (Table 2) likely acts as a mechanism to ensure their preferential translation among the various mRNAs within cells at an organism-wide level, perhaps independent of environmental or tissue-specific fluctuations in tRNA levels.

Collectively, our data on codons with Nonopt-codon_↑tRNAs_ status add to the growing support for a mechanism wherein non-optimal or rare codons, combined with elevated tRNA abundances, significantly shape translational regulation in eukaryotes.^34,44,45,47^ Further study in *T. castaneum*, possibly including assessments of protein abundance of genes with elevated usage of codons with Nonopt-codon_↑tRNAs_ status and with high usage of non-optimal codons with rare tRNA genes, will help unravel the relationships between non-optimal codon usage and translation. In addition, *in vivo* quantification of the tRNA populations in diverse tissue types in this beetle species,^37,39,40,44,77^ will help affirm whether these codons consistently exhibit high tRNA abundances, which could promote their preferential translation at an organism-wide level.

### 3.5. *T. castaneum* codon usage bias in context

Selection (s) on optimal codons with abundant tRNAs (defined here for those codons with Opt-codon_↑tRNA_ status, and typically denoted as under translational selection), may be influenced by factors such as effective population size (Ne) and genome size. In previous studies, smaller Ne (or NeS~1; and/or shorter generation times)^85,86^ or larger genomes in eukaryotes have been linked to reduced selection pressures on codon use.^87^ For example, using a statistic aimed to quantify an organism’s genome-wide selection on codon usage (using predicted selection pressures per codon and tRNAs; see also^74^) to compare among species, it was reported that strong selection pressures on codon use occurs for some bacteria such as *E. coli* (which also have highly skewed RSCU values (>2) for some of its codons^88^), with intermediate pressure in *D. melanogaster* and weak/absent in pressure humans. The authors of that study suggested that this pattern was related to their (respectively increasing) genome sizes.^87^ In this context, the overall translational selection pressures on optimal codon in *T. castaneum* (Table 1) may be expected to be moderate, and similar to those of its fellow insect *D. melanogaster* (genome sizes of 160 and 175 MB respectively).^53,89^ However, such between-taxon differences on selected codon bias could also reflect weaker pressure in smaller effective population sizes, which decrease respectively from bacteria, insects and humans.^85^ Nonetheless, our present study is largely focused on the dynamics of the most highly transcribed (top 5%) genes in the genome in *T. castaneum* (rather than all genes; Table 1), and includes analyses of not only of the translational selection on optimal codons *per se* (Opt-codon_↑tRNAs_ status), but also putative selection favouring roles of wobble and non-optimal codons (Opt-codon_wobble_ and Nonopt-codon_↑tRNAs_ status) and their relationships to tRNAs in shaping translational processes in this taxon. While our data suggest selection has been a factor in shaping the frequency of each of these types of codons in highly transcribed genes in *T. castaneum*, further similar studies in more multicellular organisms, including additional *Tribolium* species, will ascertain the breadth of such patterns across diverse metazoans.

### 3.6. Comparison of Present Optimal Codon List to a Prior Report

On a final note, it is worthwhile to mention here that the optimal codon list we present in Table 1 differs from that previously reported in *T. castaneum.*^18^ The previous report used a correlation method to determine optimal codons, and a comparison of the present primary optimal codon list in Table 1 (for the whole organism analyses) to those earlier findings is shown in Supplementary Table S2. We found that only nine of the 18 primary optimal codons identified herein, were also identified as optimal by the previous study under the correlation method^18^, even when we used very loose criteria for defining a match to that prior assessment (that is, considering all optimal codons that were defined at any level under the correlation method, regardless of whether they were the primary, secondary, or tertiary optimal codon^18^, as a match to our primary optimal codon). It has been previously argued that the use of a correlation approach can often yield a misleading list of optimal codons.^59^ Further, the R values observed for the codons defined as optimal using the prior correlation method were typically <0.1 (the highest value was 0.237, Supplementary Table S2)^18^. A range of such low values, even when statistically significant, is sometimes considered a very weak or absent correlation (R<0.3),^72,73^ and thus may not be conducive to revealing codons most often used in highly transcribed genes, as was the goal here. Moreover, we found that increased gene expression level (organism-wide expression) was not positively connected to the Fop when using the optimal codons (primary optimal codon defined as strongest correlation) identified under the prior correlation method.^18^ Rather, as shown in Supplementary Fig. S2, we found only mild variation in Fop among expression classes, and Fop was reduced in low and high expressed genes as compared to moderately expressed (Ranked ANOVA and Dunn’s P<0.05), trends inconsistent with a persistent connection between Fop and expression level. However, we did find a strong connection between expression level and Fop using the optimal codons identified herein (Table 1, Fig. 1A). The method of employing ΔRSCU between high and low expressed genes has repeatedly been shown effective for specifically revealing the optimal codons, defined as those preferentially used in the most highly transcribed genes in the genome,^14,15,17,21,24,59^ as was the present objective. Thus, the optimal codons defined herein are those most often used in highly transcribed genes, and were used for all our analyses (Table 1).

### 3.7. Conclusions

The present study has revealed the complex dynamics of codon usage in the multicellular beetle model system *T. castaneum*. We found that the majority of optimal codons in this animal model are shared at the organism-wide level and match tRNA with abundant gene copies, supporting the presence of species-wide translational selection for efficient and/or accurate translation. However, we also showed that a non-negligible subset of optimal codons varied among the four tissue types, suggesting a likelihood of tissue- and sex-specific tRNA populations, and thus localized translational selection. Based on codon optimality status and tRNA gene copies, we propose two hypotheses. The first hypothesis suggests that the usage of codons with Opt-codon_wobble_ status in highly transcribed genes in this beetle has evolved as a mechanism that slows translation, which could increase precision of translation and/or protein folding. The second hypothesis proposes that usage of codons with Nonopt-codon_↑tRNAs_ status is as a mechanism that promotes high translation of mRNA of genes with specific cellular functions, which we show here to include stress response and ribosomal protein genes.

Further study in *T. castneum*, including assessments of cellular protein levels of genes using codons with Opt-codon_wobble_ and Nonopt-codon_↑tRNAs_ status in germ line and somatic tissues, will help further unravel their potential roles in translation regulation. In addition, *in vivo* quantification of the tRNA populations in various tissue types and under stressful conditions in this beetle, as this methodology improves,^37,39,40,44,77,78^ will provide additional valuable insights into tRNA population stability and variation between tissues.

While our data suggest that the frequency of specific codons in *T. castaneum* obligately requiring wobble tRNA, similar to those non-optimal codons with few tRNAs, may be linked to translational slowing or protein-folding functions in highly transcribed genes, future follow-up studies should assess whether such codons cluster or show original use patterns at or near protein (folding) structural elements, which some research suggests may occur in certain organisms^7,9,46,90–92^, and/or whether those codons may effectively slow or pause translation.^7,9,50^ In an understudied metazoan model such as *T. castaneum,* the former may be achieved via comprehensive bioinformatics analysis of protein structural properties and codon use,^46,91^ and/or the development of a cell-free translation system allowing manipulation of codon use in mRNAs such those from *Neurospora* and *Drosophila*,^7,9^ while the latter may be informed by ribosomal profiling analyses during translation.^7,9,50^ Population-level approaches will also be valuable to further ascertaining the selection pressures acting on codon use,^86,88,93^ particularly research on the mutational spectra of codons with Opt-codon_↑tRNAs_, Opt-codon_wobble_ and Nonopt-codon_↑tRNAs_ status, to ascertain whether such codon mutations show signals of selection favouring their fixation in highly transcribed genes of *T. castaneum*.

At present, most non-traditional multicellular organisms have not had as many protocols optimized for lab-based experimental or transgenic research of codon optimization, including rates of translation elongation, protein folding, tRNA-charging, or codon-anticodon tRNA binding, as compared to the established widely studied single-celled models or *in vitro* cell lines.^7,28,34,51^ We have shown here, however, using the species *T. castaneum,* that a multifaceted approach using analyses of gene expression, tRNA genes, tissue-type, and gene functionality can be used to suggest how codon usage shapes translational optimization and regulation in a metazoan system.

## Acknowledgments

The authors thank Prof. Sue Brown at KSU for generously providing samples of *T. castaneum* for this study. This work was supported by funds from Harvard University. Thanks also to the Extavour lab members for discussions and the Bauer core sequencing facility at Harvard for generating RNA-seq data. We are grateful to the anonymous reviewers for valuable comments that helped improve our manuscript.

## SUPPLEMENTARY MATERIAL

**Table S1.**
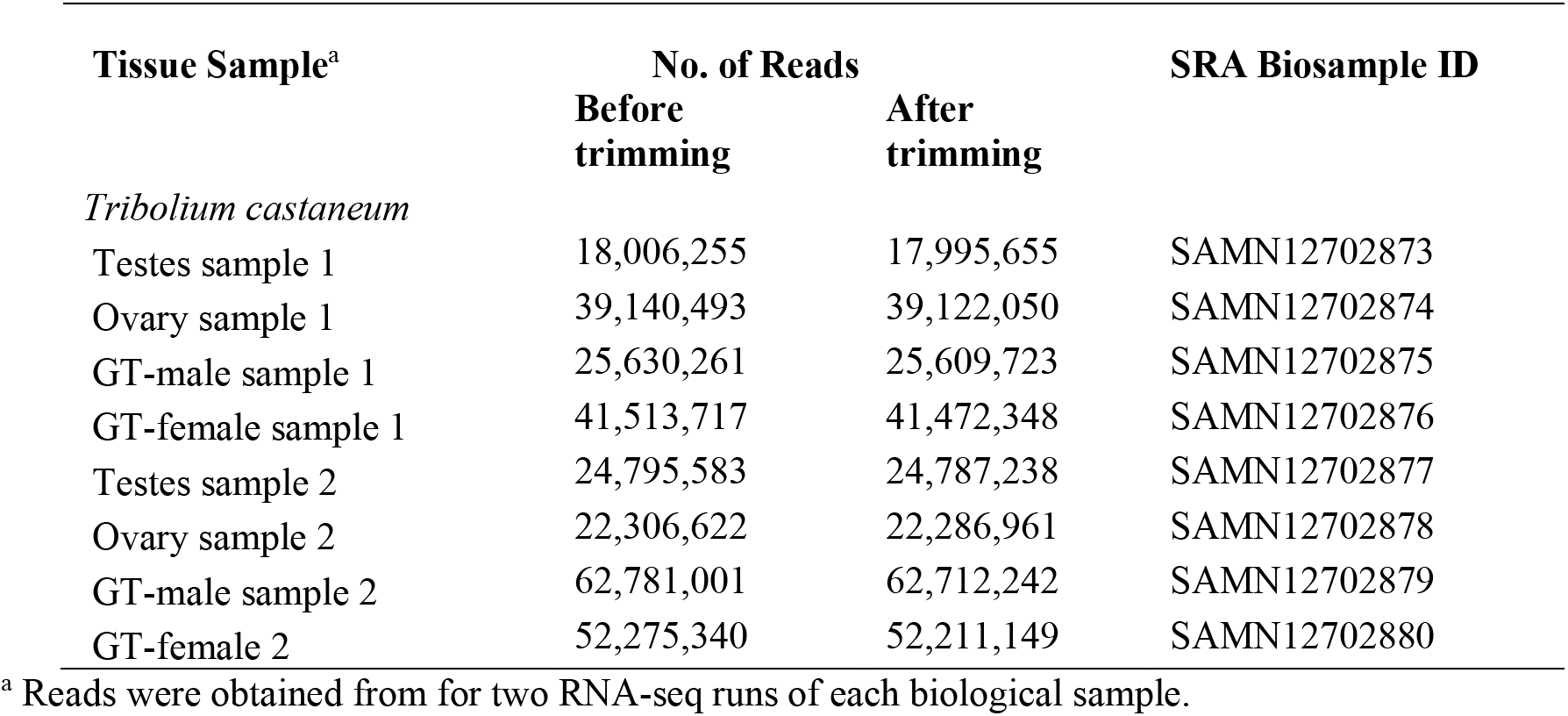
The number of RNA-seq reads for each tissue-type in the present study ^1^. RNA-seq data are shown before and after adapter and quality trimming with BBDuk (https://jgi.doe.gov/data-and-tools/bbtools/). The Short Read Archive (SRA) Biosample identifiers are also shown (https://www.ncbi.nlm.nih.gov/sra).

**Table S2.**
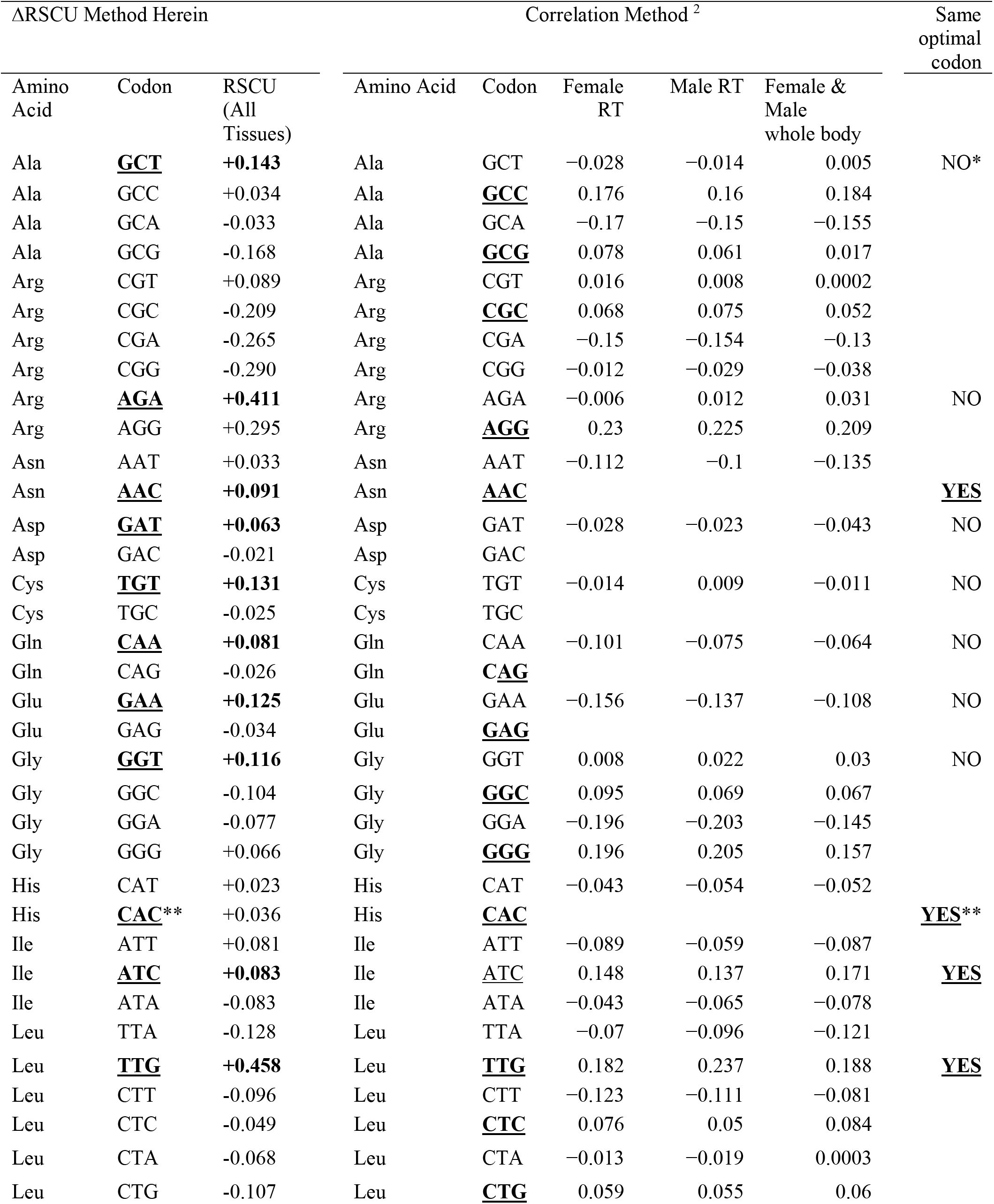
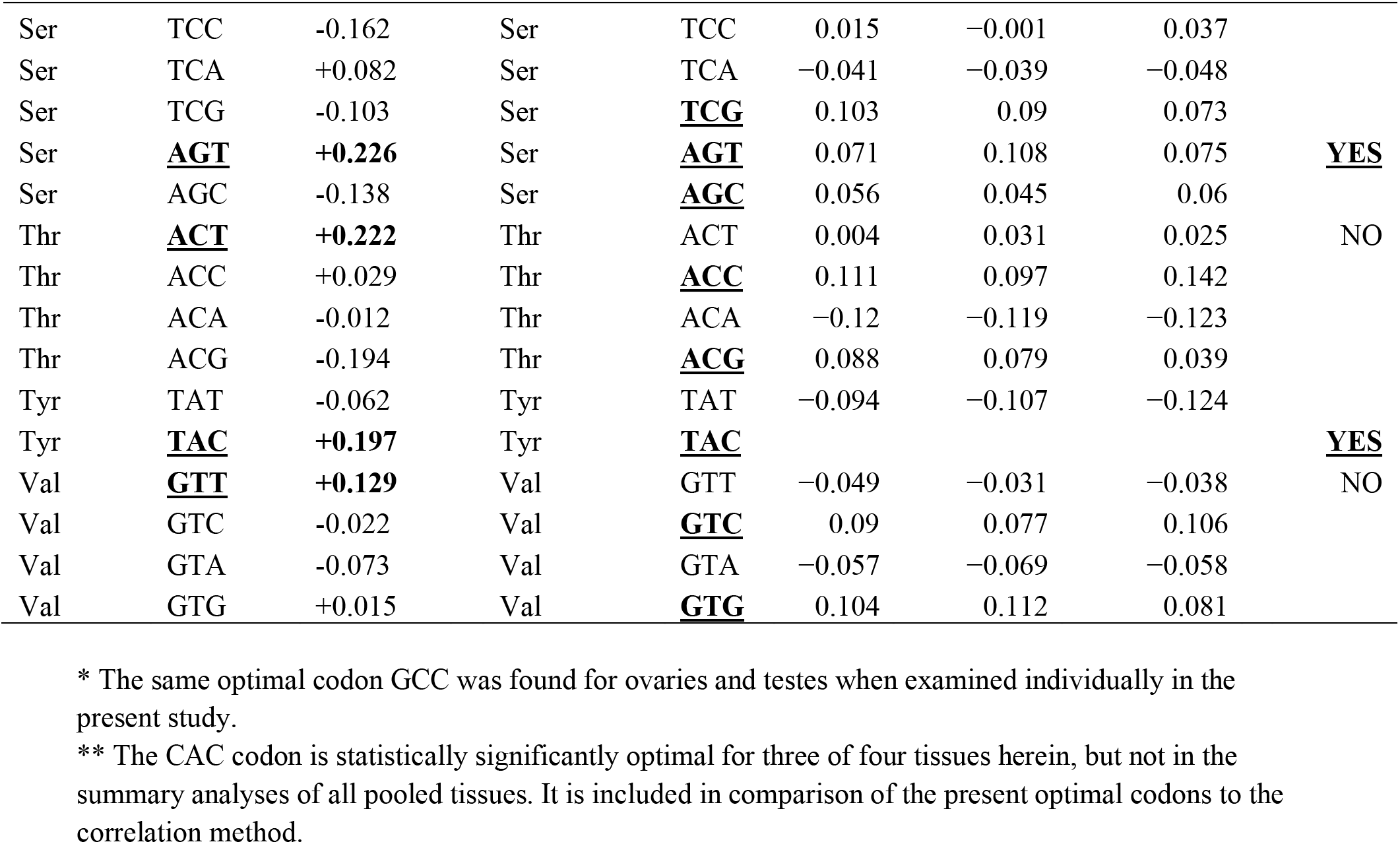
Comparison of the primary optimal codon list generated using the organism-wide analyses of high and low expressed genes in the present study (ΔRSCU) to optimal codons obtained using the correlation method in Williford and Demuth (2012), which defined up to three optimal codons per amino acid. Cases wherein the present primary optimal codon matched an optimal codon (at any level) identified under the correlation approach are indicated in the right-most column. Optimal codons defined in each study are in bold and underlined.

**Figure S1.**
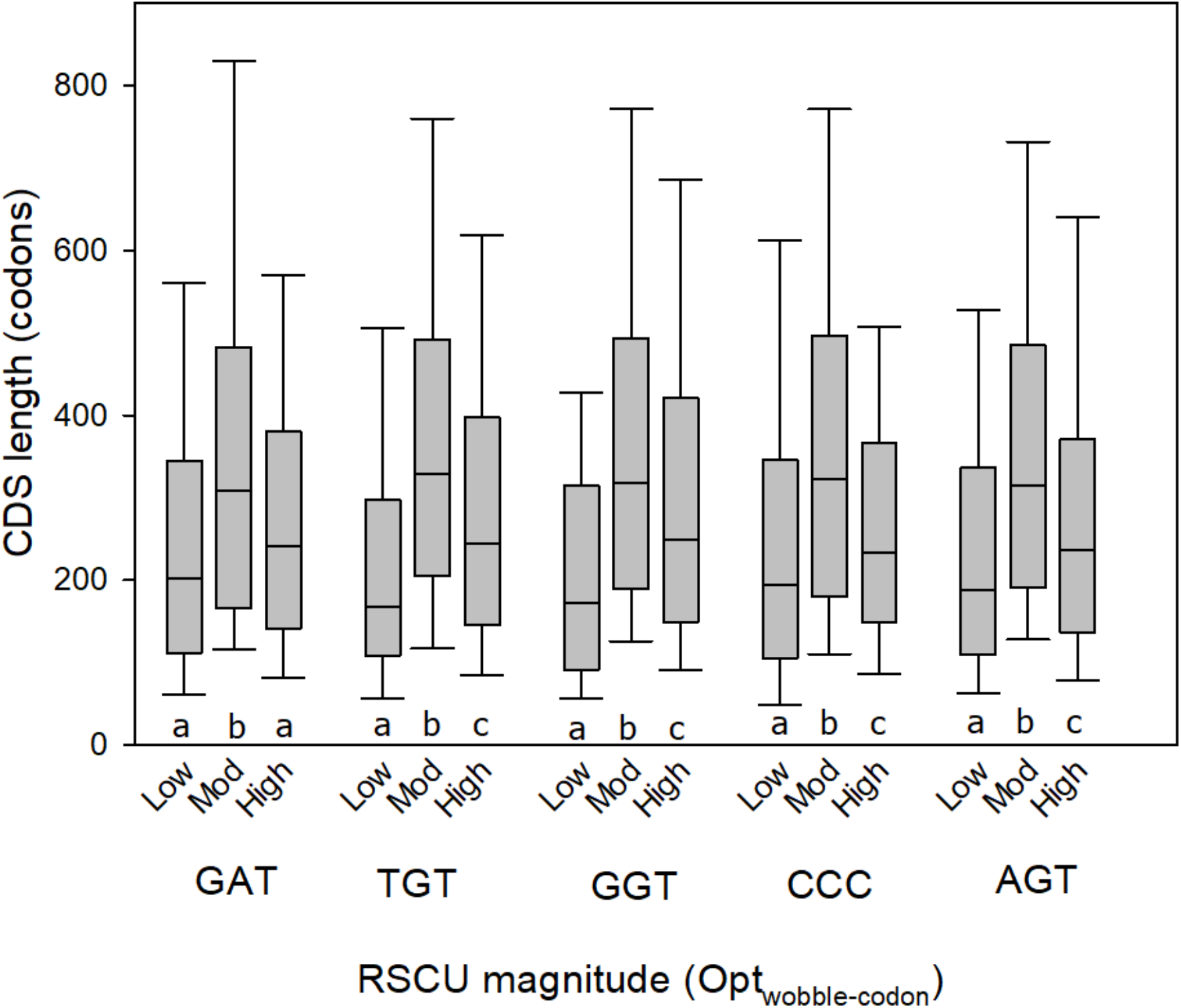
The relative use of codons with Opt-codon_wobble_ status (GAT, TGT, GGT, CCC and AGT) in highly expressed genes with respect to CDS length. Different letters below each set of three bars (per codon) indicate a statistically significant difference using Ranked ANOVA and Dunn’s paired contrasts (P<0.05). The 822 highly expressed genes were divided into three equal sized classes of RSCU values (low, moderate (mod), high) for each codon.

**Figure S2.**
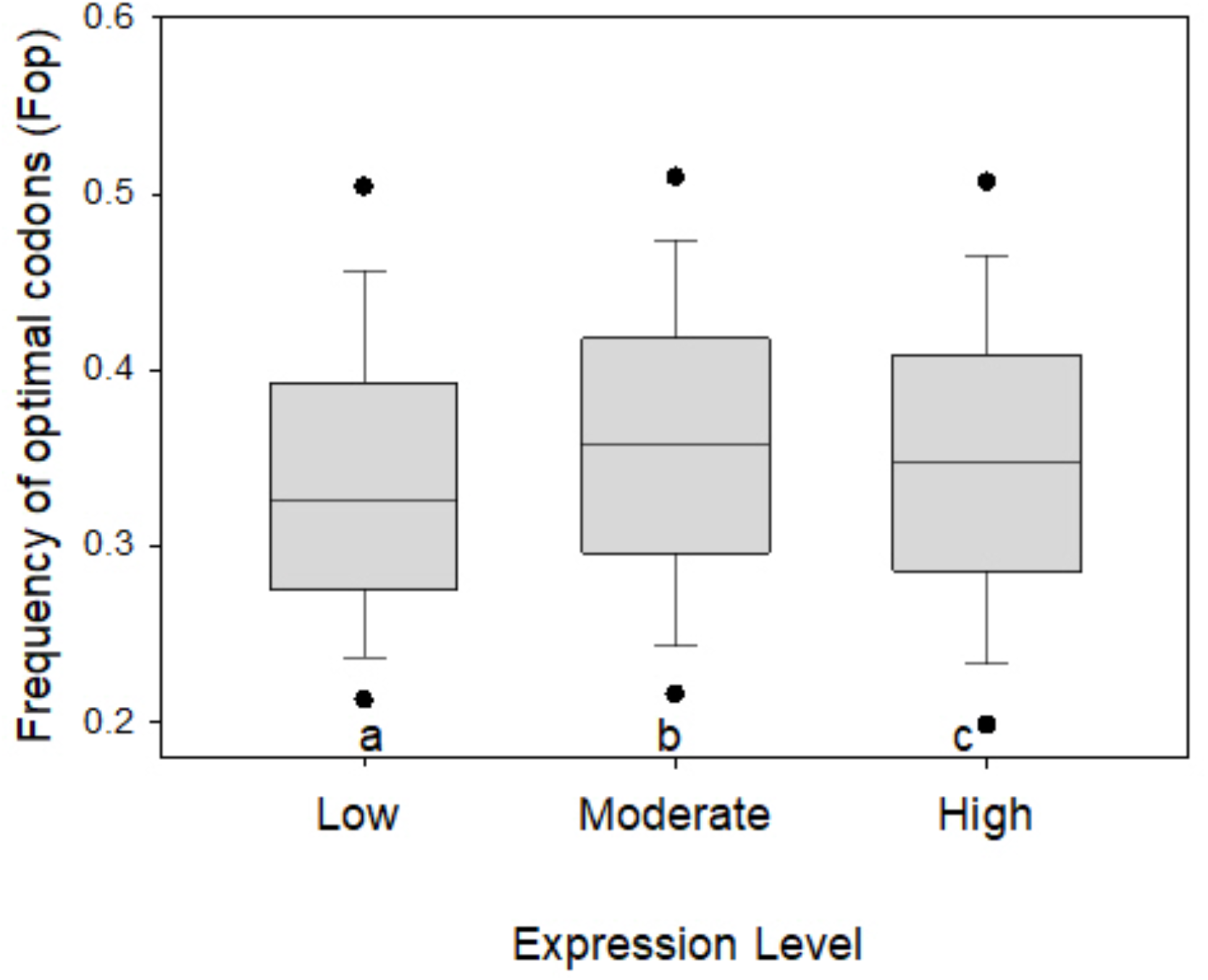
The frequency of optimal codons (Fop) across all genes studied in *T. castaneum* when using the primary optimal codons identified in Williford and Demuth.^2^ Genes are categorized into low (lowest 5%, FPKM<0.013), moderate (5 to 95%) and high (top 5%) transcription groups (FPKM>103) based on average expression across all four tissue types (testes, ovaries, GT-males, GT-females). Different letters below bars indicate a statistically significant difference using Ranked ANOVA and Dunn’s paired contrasts (<0.05).

### Supplementary Text File S1: Protein length and Opt_↑tRNA_ status

For the beetles studied herein, we found that the use of Opt-codon_wobble_ codons was connected to protein length. Specifically, for the 822 highly transcribed genes in this organism (top 5%), we ranked the RSCU for each of the five codons with Opt-codon_wobble_ status, and genes were then binned into three equal sized categories (N=274 genes each) based on the relative magnitude of RSCU (low, moderate, and high). By definition as an optimal codon, each of these five codons had elevated RSCU in the highly expressed genes (as compared to low expressed genes, Table 1). However, within the highly transcribed gene set, we found that the bin containing moderate RSCU values were consistent linked to longer CDS than those with the lowest or highest RSCU values for each of the five Opt-codon_wobble_ codons, namely GAT, TGT, GGT, CCC and AGT (Ranked ANOVA and Dunn’s paired contrast P<0.05, Fig. S1). Thus, the highly transcribed CDS encoding long proteins, appear to be connected to a specific frequency of Opt-codon_wobble_ codons, which may play a role in their translation. This may possibly comprise a mechanism to ensure a balance between high translation rates (ensured by moderate rather than highest usage of Opt-codon_wobble_ codons) and allowing intermittent pausing during translation for accurate protein synthesis and/or protein folding (ensured by their moderate, rather than low, usage) of CDS encoding long proteins.

